# NEK10 tyrosine phosphorylates p53 and controls its transcriptional activity

**DOI:** 10.1101/516971

**Authors:** Nasir Haider, Previn Dutt, Bert van de Kooij, Michael Yaffe, Vuk Stambolic

## Abstract

In response to genotoxic stress, multiple kinase signalling cascades are activated, many of them directed towards the tumour suppressor p53 which coordinates the DNA damage response (DDR). Defects in DDR pathways lead to an accumulation of mutations that can promote tumorigenesis. Emerging evidence implicates multiple members of the NimA-related kinase (NEK) family (NEK1, NEK10 and NEK11) in the DDR. Here, we describe a function for NEK10 in the regulation of p53 transcriptional activity through tyrosine phosphorylation. NEK10 loss increases cellular proliferation through modulation of the p53-dependent transcriptional output, by directly phosphorylating p53 on Y327, revealing NEK10’s unexpected substrate specificity. A p53 mutant at this site (Y327F) acts as a hypomorph, causing an attenuated p53-mediated transcriptional response. Consistently, NEK10-deficient cells display heightened sensitivity to DNA damaging agents and low *NEK10* expression is an independent predictor of a favorable response to radiation treatment in WT *TP53* breast cancer patients.

## INTRODUCTION

Eukaryotic cells respond to various forms of DNA damage through distinct DNA damage response (DDR) pathways. These molecular circuits first function to recognize damaged DNA and suppress cell cycle progression, to provide time for repair of genetic material or, if the damage is deemed irreparable, carry out an apoptotic programme. Deregulation of DDR networks leads to genetic abnormalities, chromosomal instability and accumulation of mutations that can initiate neoplastic transformation and, ultimately tumorigenesis. DDR networks are subject to complex regulation integrating multiple input signals to titer the cellular response to DNA damage. One of the master regulators of the cellular response to DNA damage is the tumour suppressor TP53.

p53 is a transcription factor that coordinates an extensive gene expression program, both basally and in response to various forms of cellular stress. TP53 transcriptional targets govern a wide range of cellular processes, including cell cycle control, apoptosis, senescence, DNA repair, metabolism, immune response and migration [1–4]. p53 binds to the p53 response elements (RE) within gene promoters, and transactivates a host of genes involved in cell cycle control, apoptosis and DNA repair, while it represses a number of pro-growth genes [1, 5, 6]. Loss of p53 can lead to genomic instability and the acquisition of oncogenic mutations that effect proliferation, transformation, therapeutic resistance and metastasis [2, 7]. *TP53* germline mutations are the cause of the hereditary disorders: Li-Fraumeni syndrome (LFS) and Li-Fraumeni-like syndromes (LFL), rare diseases that lead to a predisposition to early onset cancers [8]. Somatic mutations of *TP53* represent some of the most frequent alterations in human cancers with an incidence greater than 50% (http://p53.iarc.fr/).

The p53 protein is modulated through multiple posttranslational modifications (PTMs), including phosphorylation, ubiquitination, methylation, acetylation, and SUMOylation, in response to even low levels of DNA damage or stress, that act to direct and titer the cellular p53 response to elicit context-dependent phenotypic responses [3, 9–11]. The canonical example of this form of regulation is the MDM2-p53 axis, whereby, in healthy cycling cells, p53 protein is kept at low levels by its negative regulator, the E3 ubiquitin ligase, MDM2 and its co-regulator MDMX [12, 13]. Following DNA damage, ATM/ATR phosphorylates p53 on S15, disrupting its interaction with MDM2, leading to an accumulation of p53 protein and increased transcription of multiple genes associated with the DDR [14–18]. Further, specific PTMs such as acetylation and methylation of p53 by proteins such as Tip60/hMOF, PCAF and PRTM5, also factor into the cellular decision between growth arrest and apoptosis [19–26].

Alongside p53, multiple other inputs and signalling proteins/pathways have been implicated in the control of the cellular response to DNA damage. Recently, members of the NEK kinase family have been linked to cell cycle checkpoint control and DNA repair. In response to ionizing radiation (IR), NEK1 and NEK11 activity supports checkpoint integrity and DNA repair, whereas inactivation of NEK2 or NEK6 is required for proper checkpoint engagement [27–34]. Additionally, both NEK8 and NEK9 have been shown to participate in DNA repair and the response to replicative stress [35–37], whereas NEK10 has been found to control G2/M checkpoint integrity in response to ultraviolet (UV) irradiation [38].

Analyses of cancer genomes have correlated NEK10 status with cancer incidence and outcome. NEK10 alterations and mutations have been reported in ~2.6% of human cancers in the TCGA Pan-Cancer dataset [39–41]. Moreover, a comprehensive genome wide association study (GWAS) has identified a strong breast cancer susceptibility locus within a sub-region of human chromosome 3p24 containing *NEK10* [42], whereas lowered *NEK10* expression has been associated with poor breast cancer prognosis and higher tumour grade [38]. Despite this, little is known about the function of NEK10 in tumorigenesis and in the cellular response to clinically relevant forms of DNA damage.

In this study, we describe a function for NEK10 in the regulation of p53 transcriptional activity. Work in knockout cell lines lacking NEK10 demonstrates its involvement in the control of cellular growth, DNA replication and sensitivity to genotoxic stress. Mechanistically, we show that NEK10 regulates p53 through direct phosphorylation of Y327 within the p53 oligomerization domain and demonstrate the contribution of this phosphorylation event to the transactivation of p53-target genes. Further, genomic analyses of breast cancer patient data reveals NEK10 as a candidate prognostic indicator in WT TP53 tumours.

## RESULTS

### Loss of *NEK10* leads to increased proliferation and DNA replication

To directly investigate the biological function of NEK10, we generated A549 lung adenocarcinoma cell lines with a loss of *NEK10* function by CRISPR-Cas9 mediated deletion of exon 24 of the *NEK10* gene (A549 *NEK10^Δ/Δ^* cells). The targeted exon contains the “DFG” motif which is required for NEK10 protein kinase activity (Figure S1). Phenotypic characterization of *NEK10^Δ/Δ^* cells, revealed an increase in proliferation and colony-forming ability compared to both parental A549 cells and to *NEK10^+/+^* single cell clones (Figure 1a-c), suggesting a growth-suppressive function for NEK10.

**Figure 1.**
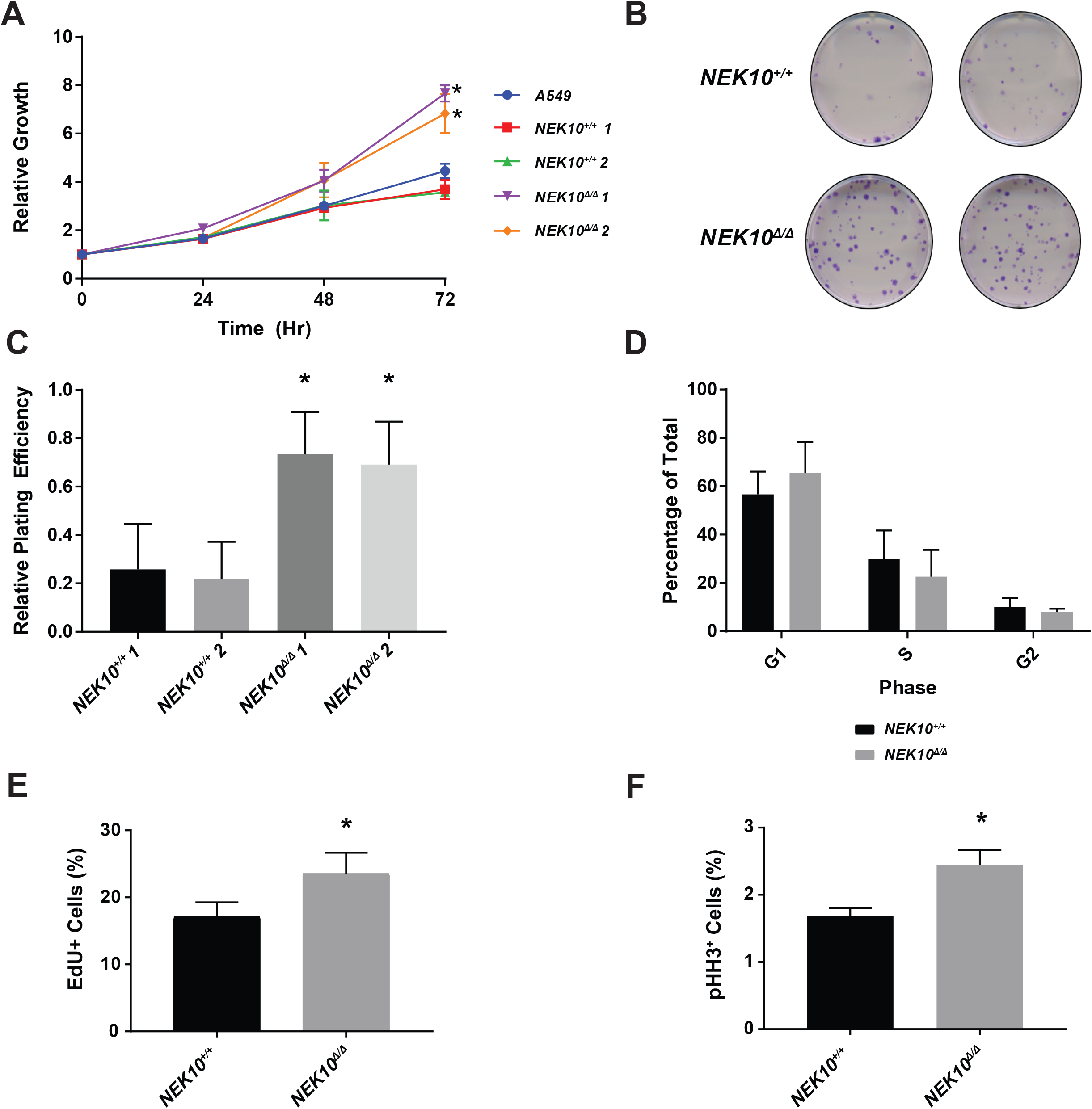
*NEK10* loss leads to an increase in cellular proliferation and DNA replication. **A**) Proliferation curve of A549 *NEK10^+/+^* and A549 *NEK10^Δ/Δ^* cells measured using SRB assay over a 72-hour time period. Relative growth was calculated relative to the absorbance at t=0 for each cell line (p<0.002, t test, n=3, bars represent SEM). **B-C**) Colony forming assay of *NEK10^+/+^* and *NEK10^Δ/È^* A549 cells. 100 cells of each genotype were seeded and their plating efficiency determined (p<0.05, t test, n=3, bars represent SEM). **D**) Cell cycle distribution was determined using propidium iodide staining for total DNA content. G1, S and G2 fractions were distinguished using FlowJo. **E**) *NEK10^+/+^* and *NEK10^Δ/Δ^* A549 cells were pulse labelled with EdU for 2 hours. Flow cytometry was performed to quantify the proportion of cells with active DNA replication. **F**) Asynchronous cells were stained with a phospho-H3 (Ser10) antibody to determine the proportion of mitotic cells. (p<0.01, t test, n=3, bars represent SEM).

Despite the differences in cellular proliferation, cell cycle distributions based on propidium iodide (PI) staining of *NEK10^+/+^* and *NEK10^Δ/Δ^* cells were indistinguishable, pointing to intact basal checkpoint fidelity in the absence of *NEK10* (Figure 1d). Considering that PI staining of asynchronous cells only provides a “snapshot” of the DNA content of a population of cells, we pulse-labeled cells with 5-ethynyl-2’-deoxyuridine (EdU), a nucleoside analog of thymidine that is incorporated into the DNA during active DNA synthesis. Measurement of the proportion of cells with active DNA synthesis (EdU+ cells), showed increased DNA synthesis in *NEK10^Δ/Δ^* cells (Figure 1e). In addition, staining of cells for phosphorylated Histone H3 (Ser10), a mitotic marker, indicated that a higher proportion of *NEK10^Δ/Δ^* cells were undergoing mitosis, further supporting a function for *NEK10* in the control of cellular proliferation (Figure 1f).

### NEK10 regulates transcription of p53 target genes

To explore the mechanism of NEK10-mediated control of cellular proliferation and DNA synthesis, we first sought to determine the activity of the pro-growth signalling PI3K and MAPK pathways and found that there was no measureable difference in the degree of pAKT and pERK levels between the *NEK10^+/+^* cells and *NEK10^Δ/Δ^* cells (Figure 2a) [43]. We next assessed the cellular levels of p53 and the p53-responsive genes p21 and MDM2 [44–47]. When compared to parental A549 and *NEK10^+/+^* cells, *NEK10^Δ/Δ^* cells displayed lowered expression of p21 and MDM2 (Figure 2b). This was not due to alterations in p21 or p53 stability, as their half-lives were indistinguishable between wildtype and *NEK10* knockout cell lines (Figure 2c). We next explored the mRNA levels of multiple p53 target genes (p21, p53, MDM2, GADD45α, PUMA, and BAX). Quantitative reverse transcription PCR (RT-qPCR) analysis revealed decreased mRNA expression of the p53-responsive genes p21, p53 and MDM2 in *NEK10^Δ/Δ^* cells, but not GADD45α, BAX, and PUMA (Figure 2d). Consistent with this observation, re-expression of NEK10 in *NEK10^Δ/Δ^* cells led to an increase in the expression of both p21 and MDM2, while leaving p53 levels unaffected, indicating a relationship between NEK10 and p53 transcriptional activity (Figure 2e).

**Figure 2.**
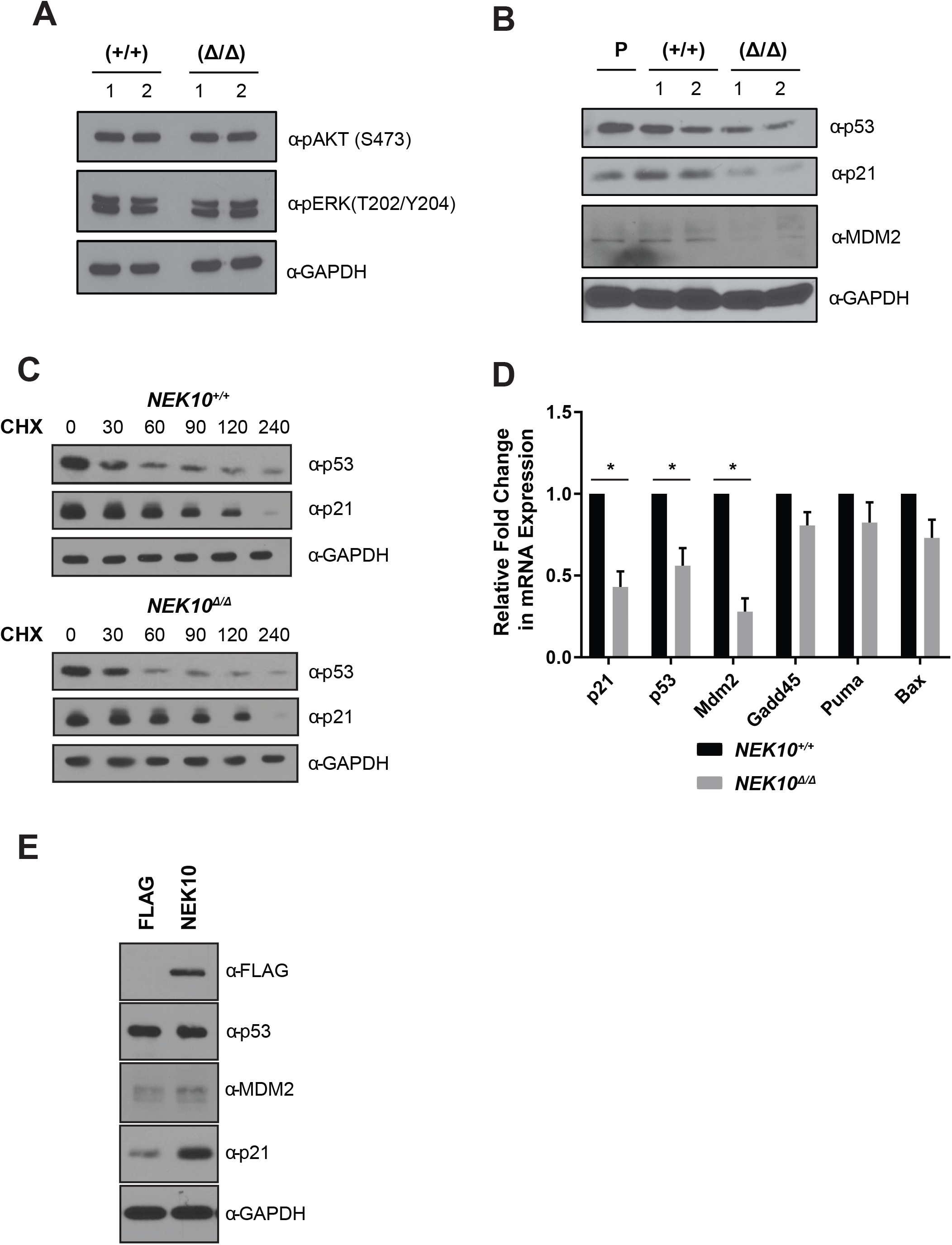
*NEK10* status modulates the expression of p53-responsive genes. **A)-B**) Immunoblot of A549 *NEK10^+/+^* and *NEK10^Δ/Δ^* cells for the expression of the indicated proteins. **C**) Cyclohexamide pulse chase experiment to determine half-life of indicated proteins. **D**) qRT-PCR of p53-responsive genes in *NEK10^+/+^* and *NEK10^Δ/Δ^* A549 cells (p<0.01, t test, n=6, bars represent SEM). **E**) Immunoblot of *NEK10^Δ/Δ^* A549 cells reconstituted with indicated lentiviral constructs.

### NEK10 phosphorylates p53 on tyrosine 327

To determine the mechanism of p53 regulation by NEK10, *NEK10^Δ/Δ^* cells were stably reconstituted with WT NEK10, a kinase activity-dead mutant NEK10 D655N or a novel serine-restricted/tyrosine kinase activity-dead mutant NEK10 I693P (Figure S2 and described in the accompanying manuscript van de Kooij et al.). While overexpression of WT NEK10 led to a strong decrease in cellular proliferation (Figure 3a), expression of either NEK10 D655N or NEK10 I693P led to a considerably less growth suppression, supporting the notion that NEK10 kinase activity, and specifically its tyrosine kinase activity, was responsible for control of proliferation. Consistent with this, only the cells expressing WT NEK10 displayed elevated expression of p21 and MDM2 (Figure 3b-c).

**Figure 3.**
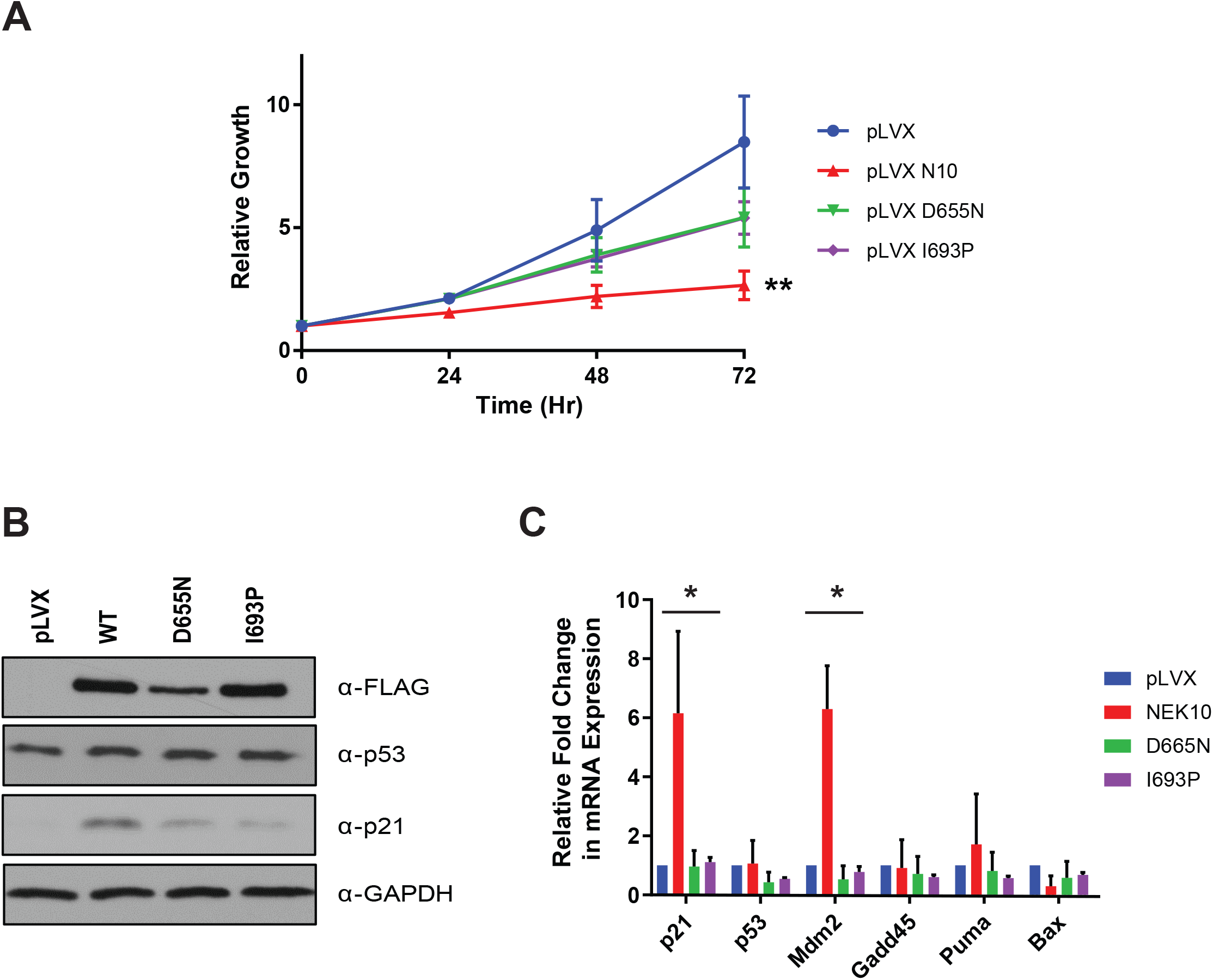
NEK10 tyrosine kinase activity is required for expression of p53-responsive genes. **A**). Proliferation measurement of *NEK10^Δ/Δ^* A549 cells reconstituted with the indicated constructs (p<0.01, t test, n=3, bars represent SEM). **B**) Immunoblot of *NEK10^Δ/Δ^* A549 cells reconstituted with the indicated constructs. **C**) qRT-PCR of p53-responsive genes in *NEK10^Δ/Δ^* A549 cells reconstituted with the indicated constructs. mRNA expression levels are expressed relative to the expression levels in pLVX-NEK10^*Δ/Δ*^ A549 cells (p<0.05, t test, n=3, bars represent SEM).

Given the observed contribution of NEK10 kinase activity to the expression of p53 target genes and growth suppression, we sought to determine if NEK10 can directly phosphorylate p53. In *in vitro* kinase assays, FLAG-NEK10 purified from HEK293T cells readily phosphorylated GST-p53 (Figure 4a). Further, overexpression of NEK10 in HEK293T cells led to an increase in tyrosine phosphorylated p53, indicating the potential site of phosphorylation is a tyrosine residue (Figure 4b). We next queried the Phosphosite™ database for reported sites of p53 tyrosine phosphorylation found in phospho-mass spectroscopic studies and identified 3 sites within p53 as candidate NEK10 target sites: Y126, Y220 and Y327 [48]. As both Y126 and Y220 had been previously characterized as sites of Src-mediated phosphorylation [48–51], we focused on Y327 as a candidate site for NEK10 phosphorylation (Figure 4c). Interestingly, Y327 also fits within a recently mapped phosphorylation site motif for NEK10, with an aromatic phenylalanine at the P+1 position (described in the accompanying manuscript van de Kooij et al.). NEK10 phosphorylated a peptide encoding p53 amino acids 320-335, but failed to phosphorylate the same peptide with a Y327F substitution (Figure 4d-e). Despite the peptide containing two candidate phospho-acceptor sites (Y327 and T329), phosphorylation of the peptide appeared to be wholly dependent on Y327 as a phospho-acceptor residue, as NEK10 readily phosphorylated the T329A mutant peptide. Moreover, NEK10 failed to phosphorylate full length Y327F GST-p53, reinforcing Y327 as a site for NEK10 phosphorylation (Figure 4f-g). Consistent with an active function of NEK10 in p53 Y327 phosphorylation, WT p53 was tyrosine phosphorylated in *NEK10^+/+^* but not *NEK10^Δ/Δ^* cells, whereas the p53 Y327F mutant was not tyrosine phosphorylated in either of the cell lines (Figure 4h).

**Figure 4.**
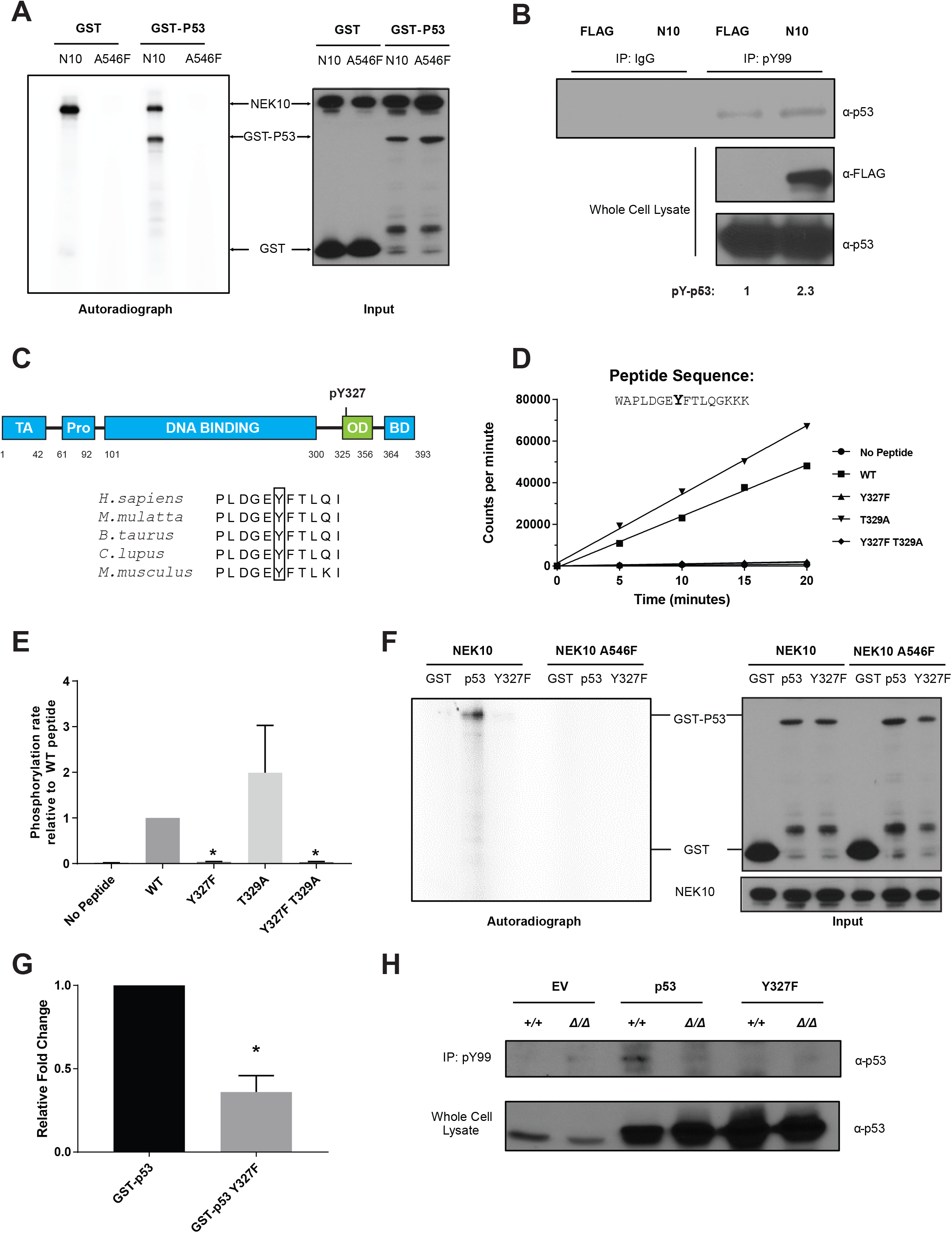
NEK10 phosphorylates p53 on Y327 *in vitro*. **A**) Radioactive *in vitro* kinase assay performed using purified FLAG-NEK10 WT and kinase dead FLAG-NEK10 (A546F) incubated with GST alone or GST-p53. **B**) HEK293T cells transfected with the indicated constructs were lysed under denaturing conditions and tyrosine phosphorylated proteins were immunoprecipitated. The amount of pY-p53 was determined by anti-p53 immunoblotting. **C**) Schematic representation of p53 domain organization highlighting the location of Y327 with the relevant domains indicated. **D**) Radioactive *in vitro* kinase assay performed using purified FLAG-NEK10 WT and a peptide centered on Y327. Incorporated radioactive ^32^P was quantified via liquid scintillation at the indicated time points. **E**) Quantification of relative rates of phosphate transfer for WT peptide and phosphoacceptor site mutants (p<0.05, t test, n=3, bars represent SEM). F) Radioactive *in vitro* kinase assay performed using purified FLAG-NEK10 WT and the indicated GST-p53 constructs. **G**) Quantification of relative phosphate transfer (p<0.05, t test, n=3, bars represent SEM). **H**) Indicated p53 constructs were transfected into *NEK10^+/+^* and *NEK10^Δ/Δ^* A549 cells and the phosphotyrosine-containing proteins were immnoprecipitated. The amount of pY-p53 was determined by anti-p53 immunoblotting.

### NEK10 phosphorylation of p53 tyrosine 327 supports p53 transcriptional activity

We next investigated the function of Y327 in p53 transcriptional activity, by transfecting HCT116 *p53^−/−^* cells with WT p53, p53 Y327F, and p53 L344P. Judging by transactivation of a panel of p53-target genes and the protein levels of both p21 and MDM2, p53 Y327F was not a full loss-of-function p53 mutant but rather acted as a hypomorph to WT p53 (Figure 5a-c). Y327F substitution impaired p53 transcriptional activity, but to a lesser extent than the one caused by the L344P mutation, which leads to complete loss of p53 ability to influence transcription (Figure 5a-c) [52]. Even though Y327 localizes to the oligomerization domain of p53, and previously reported data indicating that nitration of this amino acid influenced oligomerization and transcriptional activity, Y327F mutation failed to impact p53 oligomerization in our system (Figure S3) [53, 54].

**Figure 5.**
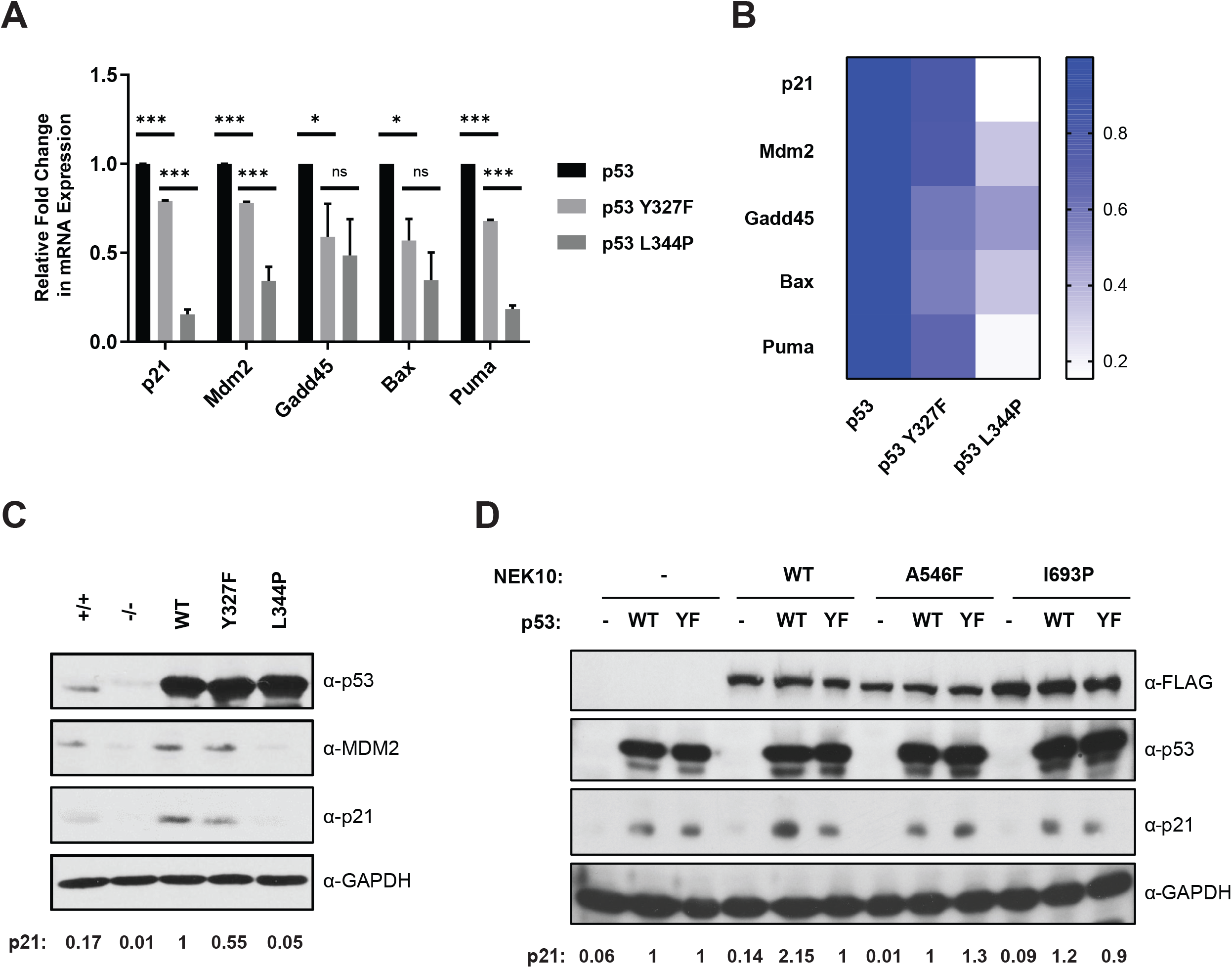
Mutation of p53-Y327 attenuates the expression of p53 target genes. **A**) qRT-PCR of p53 responsive genes in HCT116 *p53^−/−^* cells reconstituted with indicated constructs to determine relative expression levels compared to WT-p53 (*= p<0.05, **=p<0.01, ***=p<0.001, t test, n=3, bars represent SEM). **B**) Heatmap of qRT-PCR of p53 responsive genes **C**) Immunoblot of HCT116 *p53^−/−^* cells reconstituted with indicated His-p53 constructs. Quantifications of the expression levels of p21 relative to the cells overexpressing WT-p53 are indicated. **D**) Immunoblot of H1299 cells reconstituted with the indicated variants of FLAG-NEK10 and His-p53. Quantifications of expression levels of p21 relative to cells overexpressing both FLAG and WT-p53 are indicated.

The relationship between p53 Y327 and NEK10 kinase activity was further assessed by co-expression of WT, kinase dead (A546F), or serine-restricted/tyrosine kinase activity-dead (I693P) NEK10 with WT p53 or p53 Y327F, respectively, in H1299 cells (p53-null) (Figure 5d). p21 protein levels were only increased by the co-expression of WT p53 and WT NEK10, pointing to NEK10 as an inducer of p53 transcriptional activity. This effect was highly dependent on the intact Y327, as the increase in p21 levels diminished in cells co-expressing WT NEK10 and p53 Y327F. Accordingly, kinase-deficient mutants of NEK10 (A546F, I693P) failed to affect p21 levels in combination with WT p53 (Figure 5d).

### Induction of p53 target genes in response to genotoxic stress requires NEK10

In light of NEK10’s impact on p53 transcriptional activity and previous work implicating NEK10 in the DNA damage response [38], we next investigated the effect of NEK10 loss in the context of DNA damage. In response to cisplatin treatment, NEK10 loss impaired induction of certain p53 responsive genes, such as p21, GADD45α, and PUMA (Figure 6a). This effect was independent of p53 S15 phosphorylation, or the fluctuations in p53 protein stability, as these remained indistinguishable between *NEK10^+/+^* and *NEK10^Δ/Δ^* cells (Figure 6a). This defect in p53 activity was also observable in response to IR, as irradiation led to a rapid and sustained increase in both p21 and p53 protein levels in the *NEK10^+/+^* cells, reaching maximum levels 30 minutes after exposure, in contrast to the *NEK10^Δ/Δ^* cells which took up to 4 hours to build up maximum levels of p21 expression (Figure 6b, c). Exposure of cells to IR also prompted tyrosine phosphorylation of WT p53, which was abolished by p53 Y327F substitution (Figure 6d). Supporting the importance of Y327 phosphorylation for the p53 transcriptional response, reconstitution of H1299 cells (p53-null) with the p53 Y327F mutant failed to fully support expression of p21 and MDM2 following IR, when compared to the same cells expressing WT p53 (Figure 6e).

**Figure 6.**
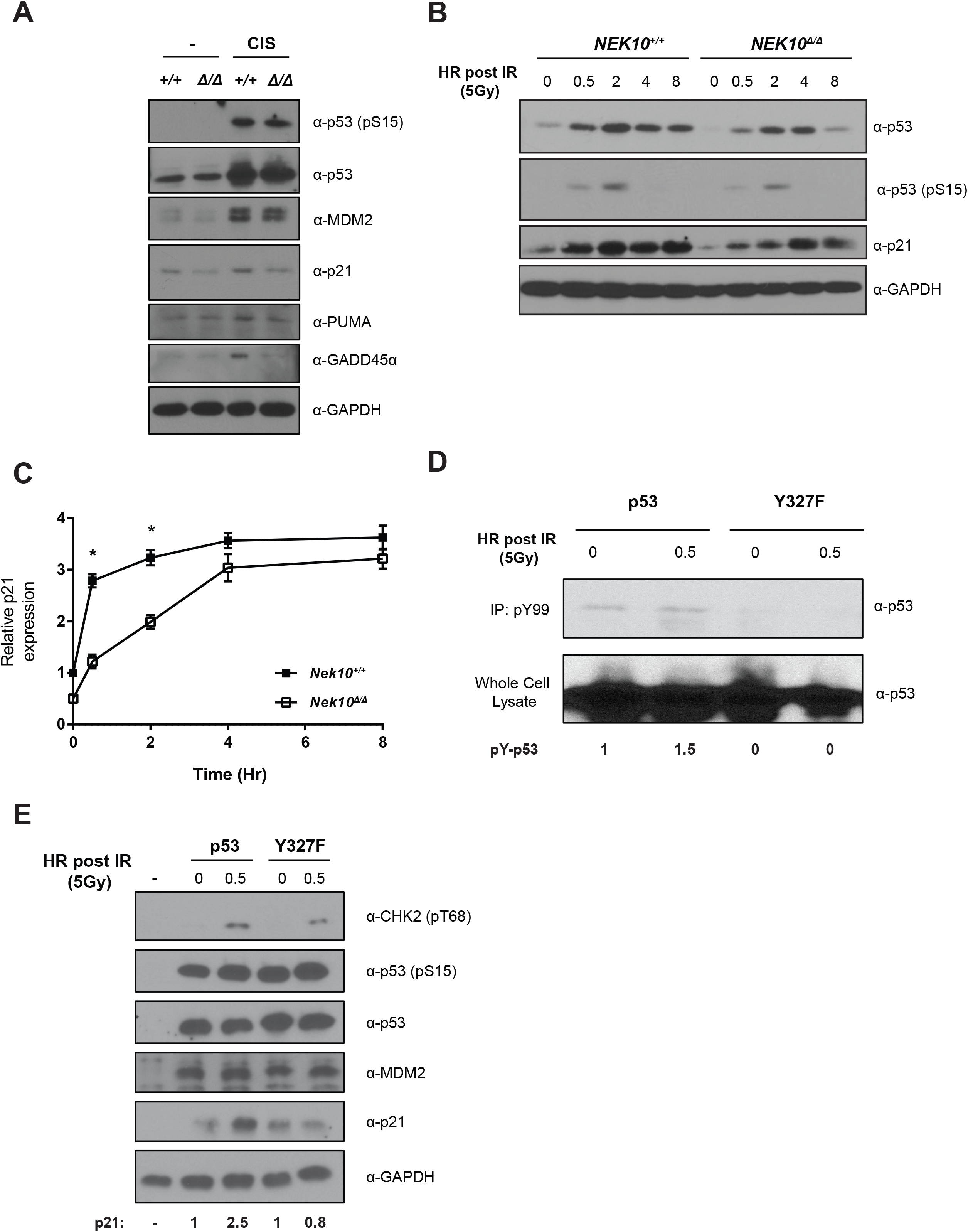
*NEK10* modulates the expression of p53 responsive genes is response to genotoxic agents. **A**) *NEK10^+/+^* and *NEK10^Δ/Δ^* A549 cells were exposed to 5 μM of cisplatin and were lysed 20 hours after treatment **B**) *NEK10^+/+^* and *NEK10^Δ/Δ^* A549 cells were exposed to 5 Gy of IR and were lysed at the indicated time points after treatment. **C**) Quantifications of the expression levels of p21 relative to the p21 levels of *NEK10^+/+^* cells at T0 (n=3, bars represent SEM, *=p<0.005). **D**) H1299 cells transfected with the indicated constructs were treated with 5 Gy of ionizing radiation and lysed at the indicated timepoints under denaturing conditions and tyrosine phosphorylated proteins were immnoprecipitated. The amount of pY-p53 was determined by anti-p53 immunoblotting. **E**) H1299 cells transfected with the indicated constructs were lysed under denaturing conditions in response to IR and immunoblots were performed to monitor the expression of the indicated proteins. Quantifications of the expression levels of p21 relative to the untreated cells are indicated.

We next probed the effect of *NEK10* status on cellular sensitivity to escalating doses of genotoxic agents using clonogenic assays. *NEK10* loss led to increased cell sensitivity to cisplatin and olaparib (Figure 7a-b). This effect was paralleled by the impact of *NEK10* loss on the DNA damage-induced cell cycle arrest. In response to cisplatin treatment, and in contrast to *NEK10^+/+^* cells, *NEK10^Δ/Δ^* cells were compromised in the induction of the G2/M arrest (Figure 7c-d). Compared to *NEK10^+/+^* cells, G2/M arrest defect in *NEK10^Δ/Δ^* cells was accompanied by heightened cell death, evidenced by the increased proportion of both the sub-G1 and Annexin V+/PI+ cells, (Figure 7e-f).

**Figure 7.**
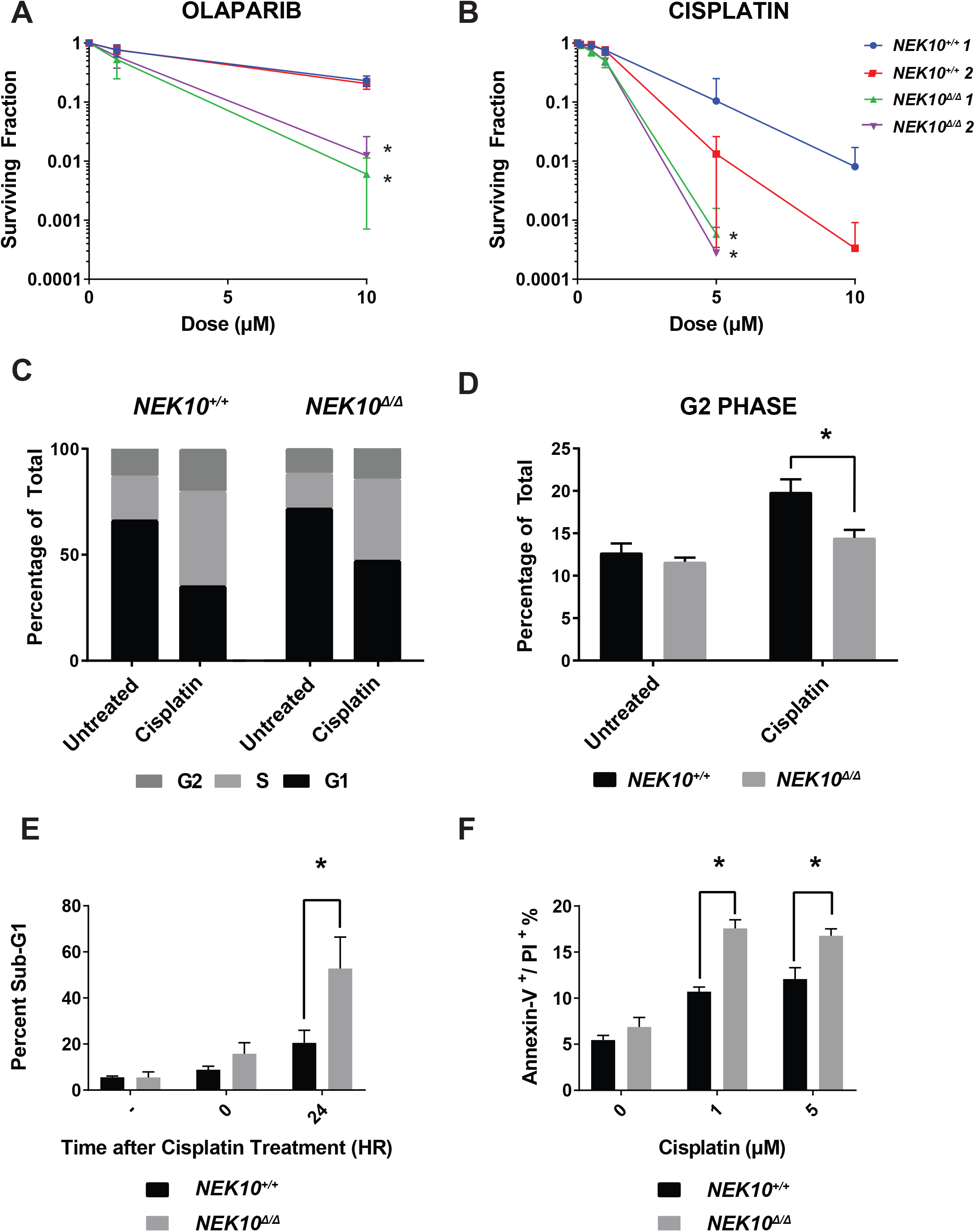
*NEK10* loss sensitizes cells to chemotherapeutic agents. **A)-B**) *NEK10^+/+^* and *NEK10^Δ/Δ^* A549 cells were treated with the increasing doses of olaparib or cisplatin and assessed for clonogenic survival 10-12 days after treatment. Graph represents percent of cells that survive after treatment compared to untreated control (p<0.005, t test, n=3, bars represent SEM). **C**) Cells were treated with 5 μM of cisplatin overnight and allowed to recover for 24 H prior to being analysed for DNA content with propidium iodide. **D**) Quantification of G2 populations for cisplatin treated cells as an indicator of G2/M arrest (p<0.01, t test, n=4, bars represent SEM). **E**) Cells were treated with 5 μM cisplatin for 20 hours and then assessed for proportion of sub-G1 cells, an indicator of cell death, at the indicated time points after treatment (p<0.05, t test, n=3, bars represent SEM). **F**) Cells were treated with indicated doses of cisplatin and assessed for apoptosis with Annexin V/propidium iodide staining 24 hours after overnight cisplatin treatment.

### Reduced *NEK10* expression is associated with poor prognosis and increased radiosensitivity in breast cancer patients

In light of previous evidence of genetic alterations in and around the *NEK10* gene in cancer [38–42, 55, 56], and the function in p53 regulation we uncovered, we sought to determine if *NEK10* mRNA expression related to clinical outcome in cancer. Querying The Cancer Genome Atlas (TCGA), we found that reduced *NEK10* mRNA expression correlates with a significant decrease in 10-year survival in breast cancer patients (High *NEK10* vs Low *NEK10)* (p=0.0088) (Figure 8a). In agreement with our experimental data, this correlation was particularly evident when *TP53* status was taken into account, as reduced *NEK10* expression was associated with a 16% reduction in 10-year survival only in patients whose tumours were WT for *TP53* (p=0.0007) (Figure 8b, left panel), whereas no correlation was evident in patients with somatic *TP53* mutations (MUT *TP53*) (Figure 8b, right panel).

**Figure 8.**
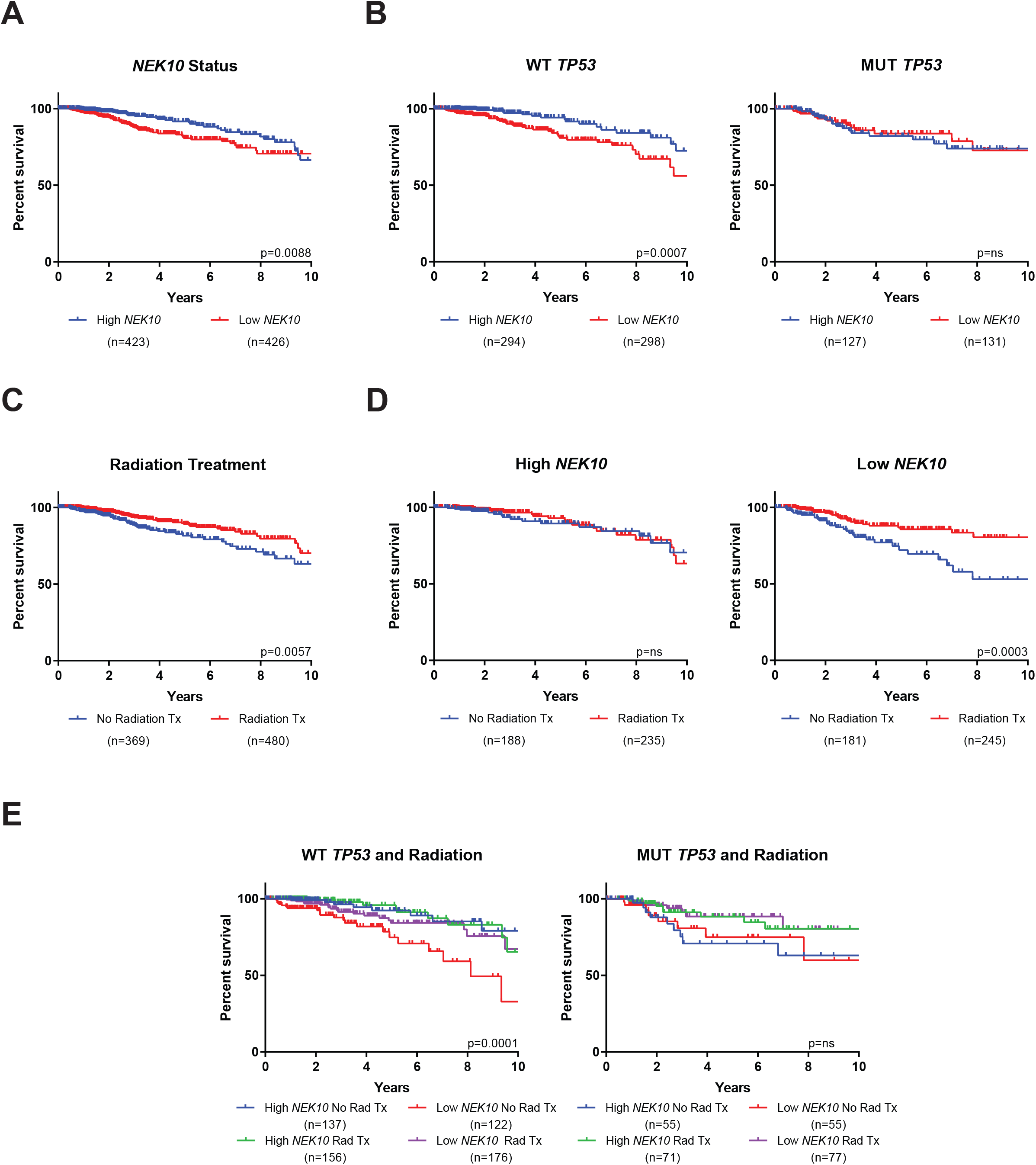
Reduced *NEK10* expression correlates with poor patient survival. **A**) Kaplan-Meier curve showing overall survival analysis in patients with high and low expression of *NEK10* mRNA from TCGA breast cancer dataset, with patients dichotomized into High *NEK10* and Low *NEK10* groups based on the median mRNA expression of *NEK10* **B**) Kaplan-Meier curve showing overall survival in patients with either WT *TP53* or MUT *TP53* relative to *NEK10* mRNA expression **C**) Kaplan-Meier curve showing overall survival analysis in patients that had either received radiotherapy (Radiation Tx), or did not receive radiation as part of their treatment regimen (No Radiation Tx) **D**) Kaplan-Meier curve showing overall survival in patients with either High or Low levels of *NEK10* expression in the context of radiation treatment. **E**) Kaplan-Meier curve showing overall survival in patients with either WT *TP53* or MUT *TP53* relative to *NEK10* mRNA expression in the context of radiation treatment. The p-values indicated were calculated using log rank analysis as indicated, and the number of patients in each group is also indicated.

Radiotherapy is often administered as a means to prevent breast cancer recurrence, and is associated with a moderate increase in overall survival (p=0.0057) (Figure 8c) [57, 58]. Interestingly, in the Low *NEK10* cohort, there was a 27% increase in 10-year overall survival for patients who had received radiation therapy compared to those that had not (p=0.0003) (Figure 8d, right panel). In contrast, there was no additive benefit of radiation therapy in the High *NEK10* patient subset (Figure 8d, left panel). The radio-response of tumours with WT *TP53* was highly dependent on *NEK10* expression. In particular, tumours from the Low *NEK10* cohort were the most responsive to radiotherapy, as these patients saw a 35% increase in overall survival (p=0.0001) (Figure 8e, left panel), compared to no significant effect on survival in the High *NEK10* cohort. In contrast, patients with somatic *TP53* mutations (MUT *TP53)* saw a benefit with radiation, but this benefit was independent of *NEK10* status (Figure 8e, right panel).

## DISCUSSION

The cellular response to DNA damage is a well-orchestrated sequence of molecular events within a complex network of intersecting signaling pathways. The process is modulated in a context-dependent manner and is sensitive to the phase of the cell cycle in which the damage is encountered, the type/degree of the damage, the extracellular milieu, or the mutational status of the cell itself. Central to many of the cellular damage response pathways is the tumour suppressor p53 [2, 59, 60]. The work presented here uncovers a function for the protein kinase NEK10 in regulating p53 transcriptional activity through phosphorylation of p53 Y327.

Loss of *NEK10* led to an increase in cellular proliferation and DNA replication (Figure 1). This is in contrast to the effects of deletion or knockdown of other NEK family kinases, including the close NEK10 homologues NEK6 and NEK7, whose loss of function leads to a decrease in cellular proliferation or induction of cell death [61–63]. The mechanism behind the NEK10-dependent effect on cell proliferation appeared to be related to a decrease in the expression of p53-responsive genes, key regulators of cell cycle progression (Figure 2b, d) [46, 47], as well as a decrease in p53 expression upon *NEK10* loss, possibly via the auto-regulatory p53 response element that exists in the human *TP53* promoter (Figure 2b, d) [64–66]. Of note, the pro-apoptotic p53 responsive genes, BAX and PUMA, were also decreased in expression but to a lesser degree, likely indicative of an intact apoptotic axis (Figure 2d). This is reflected in equal proportions of apoptotic cells under basal conditions of both *NEK10^+/+^* and *NEK10^Δ/Δ^* A549 cells (Figure 7e-f).

The differences in p53 target gene expression were dependent on the tyrosine kinase activity of NEK10 (Figure 3). Using a novel serine phosphorylation-restricted NEK10 mutant (I693P), we demonstrate that the NEK10 tyrosine kinase activity is responsible for p53 regulation (Figures 3, 5d) (accompanying manuscript van de Kooij et al.). Consistent with this, re-expression of the I693P mutant in *NEK10^Δ/Δ^* cells failed to augment p53 target gene expression (Figures 3b-c and 5d). Of interest, while re-expression of kinase defective variants of NEK10 (D655N and I693P) in *NEK10^Δ/Δ^* cells failed to decrease cellular proliferation to the degree that WT NEK10 had, a quantifiable decrease was observed when compared to *NEK10^Δ/Δ^* cells expressing a control vector (pLVX), which suggests a potential kinase-independent contribution of NEK10 to proliferative control (Figure 3a). Despite this, the kinase-independent function of NEK10 on cellular proliferation is p53-independent, as neither kinase-defective mutant was capable of stimulating p53 transcription (Figure 3b-c).

*In vitro* enzymatic assays and analysis of phosphoproteomic datasets using the tyrosine phosphorylation site motif of NEK10, uncovered Y327 as a candidate site of NEK10 phosphorylation (Figure 4). Judging by the of differential expression of the p53-responsive genes, p53 Y327 phosphorylation may establish a separate threshold for transactivation of subsets of p53 targets as the Y327F mutant induces an intermediate level of p53 target gene expression when compared to WT p53 and the loss of function p53 L344P mutant for the majority of genes assayed, reflective of the partial loss of p53 function in NEK10-deficient cells (Figures 2 b-c and 5a-c). Therefore, the Y327F p53 mutant represents a unique hypomorphic variant of p53.

This hypomorphic p53 response elicited by *NEK10* loss was manifested by differential reaction to DNA-damaging agents, such as cisplatin, where despite comparable increases in p53 protein and p53 S15 levels in response to this genotoxic treatment, *NEK10^Δ/Δ^* cells failed to induce expression of several p53 target genes (Figure 6a). Further, upon IR exposure, *NEK10^Δ/Δ^* cells displayed protracted kinetics of p53 and p21 induction (peaking at 8 hours) compared to a rapid induction and an early plateau in *NEK10^+/+^* cells (0.5 hours) (Figure 6b-c). The difference in induction kinetics of the p53 pathway likely accounts for the phenotypic results observed in cell survival and cell cycle arrest, discussed below [67–73]. While the changes in p53 tyrosine phosphorylation in response to genotoxic stress tracked with the previously well-characterized changes in p53 S15 phosphorylation and an increase in p53 transcriptional activity, Y327F mutation or *NEK10* loss did not affect S15 phosphorylation. (Figure 6 a-b, d-e). This observation suggests that Y327 phosphorylation does not affect S15 phosphorylation and S15-mediated p53 stabilization, but likely acts to promote transactivation of p53 target genes.

The weakened p53 response in *NEK10^Δ/Δ^* cells likely contributed to their sensitivity to olaparib and cisplatin when compared to the *NEK10^+/+^* cells, presenting as increased apoptosis and decreased clonogenic survival (Figure 7). *NEK10^Δ/Δ^* cells failed to engage an effective G2/M arrest in response to cisplatin possibly due to the lack of induction of GADD45α, a key mediator of the G2/M checkpoint, contributing to accruement of deleterious DNA lesions and subsequent apoptosis (Figure 6a, 7e-f) [74–76]. The observed NEK10-dependent differences in cell cycle arrest and apoptosis may indicate a selective response to genotoxic stressors, whereby the hypomorphic p53 response fails to elicit cell cycle arrest, yet is able to support an apoptotic response. Previous reports that the acetylation of p53 on Lys320 by PCAF leads to preferential transactivation of cell cycle arrest genes over the apoptotic target genes [23–25] supports the existence of post-translational modifications-driven fine-tuning of the p53 response.

Consistent with the diminution of p53 function in NEK10-deficient cells, analysis of cancer genome data revealed associations of lowered *NEK10* expression with worse outcome in breast cancer, further suggestive of a tumour-suppressive function of *NEK10*, particularly in patients lacking somatic *TP53* mutations (Figure 8a,b) [38–42]. In contrast to breast cancer patients receiving other therapies, those with reduced expression of *NEK10* displayed a beneficial response to radiation therapy, indicated by their grater overall survival (Figure 8d). The improvement in therapeutic response appeared to require p53, as only tumours with WT *TP53* showed the association between *NEK10* expression and radioresponse, reinforcing a p53-dependent function for *NEK10* in cancer (Figure 8e). Thus, patients with WT *TP53* and reduced *NEK10* expression represent a cohort that would likely benefit from radiotherapy, as lack of such treatment in this group led to a striking increase in cancer recurrence.

Taken together, our results demonstrate that NEK10 supports p53 transcriptional activity through Y327 phosphorylation, with loss of NEK10 causing an impaired p53 response and a hypomorphic p53 phenotype. Recent studies have identified subsets of breast cancers that are highly sensitive to genotoxic agents and radiation therapy, such as those with DDR defects or genomic instability (i.e. BRCA1/2 deficiency or “BRCAness”) [77–83]. Breast cancers with reduced NEK10 expression may constitute a similar breast cancer subset that is exquisitely sensitive to genotoxic treatments and radiotherapy due to an attenuated p53 response, rather than a direct defect in DNA repair. Thus, low *NEK10* expression levels represent a potential prognostic biomarker for the already standard of care chemo/radiotherapeutic treatments for breast cancer. Moreover, the yet to be developed NEK10 kinase inhibitors are predicted to act as chemo- and radio-sensitizing agents in breast cancer treatment and improve patient outcomes.

## MATERIALS AND METHODS

### Antibodies

The following antibodies were purchased from Santa Cruz Biotechnology: Anti-GAPDH FL-335 (sc-25778), Anti-p53 FL-393 (sc6243), Anti-p-Tyr PY-99 (sc-7020), Anti-GADD45α C-4 (sc-6580). Anti-FLAG M2 (F3165) and Anti-FLAG M2 Agarose were from Sigma-Aldrich. Anti-p21 (556431) was from BD Pharmingen. Anti-MDM2 Ab-1 (OP46) was from Calbiochem. Anti-PUMA (NB500-261) was from Novus Biologicals. Anti-p53 S15 (9284) was from Cell Signalling Technology.

### Cell culture, transfections, viral infections and reagents

HEK293T, HEK293T *NEK10^Δ/Δ^*, A549, A549 *NEK10^Δ/Δ^*, H1299. HCT116 *p53^+/+^* and HCT116 *p53^−/−^* cells and their derivatives were maintained in DMEM (Corning), supplemented with 10% FBS (Wisent) and Pen/Strep (100 mg/ml, Hyclone).

Transient transfections of HEK293T cells were performed by using the calcium phosphate method. Transfection of other cell lines was achieved using PolyJet™ according to manufacturer’s instructions.

All plasmids were constructed with the exception of the pLVX-3xFLAG-NEK10 series of constructs (Gift of Dr. Michael Yaffe). For expression in mammalian cells, NEK10 was cloned into 3xFLAG-CMV-7.1 NEK10, and p53 was cloned into pcDNA3-His. For expression of recombinant proteins in bacterial cells, p53 was cloned into pGEX2TK. All point mutants were generated by using the QuickChange site-directed mutagenesis kit (Stratagene),

A549 *NEK10^Δ/Δ^* cells were engineered to express pLVX-FLAG, FLAG-NEK10 WT, FLAG-NEK10 D665N and FLAG-NEK10 I693P via lentiviral transduction. In short, transduction was performed with lentiviral particles in the presence of protamine sulfate (5 μg/mL) for 20 hours. Subsequent to transduction, cells were reseeded into 200 μg/mL hygromycin (Sigma) and selected until negative control cells died (3-5 days).

### Cell lysis, immunoblotting and immunoprecipitations

Unless indicated otherwise, for immunoblotting, cells were lysed in Laemmli sample buffer (50 mM Tris-HCl pH 6.8, 2% SDS, 10% glycerol, 5% β-mercaptoethanol), normalized for total protein content, resolved by SDS-PAGE, and transferred to PVDF membranes (Millipore). Membranes were blocked in 5% BSA and probed with the indicated antibodies.

For immunoprecipitations, cells were lysed in RIPA buffer (50 mM Tris-HCl pH 7.4, 150 mM NaCl, 1% Triton X-100, 1% sodium deoxycholate, 0.1% SDS, 1 mM EDTA, supplemented with fresh 1 mM dithiothreitol (DTT), 0.1 mM sodium orthovanadate and a protease inhibitor cocktail (Sigma)) and sonicated followed by incubation on ice for 20 minutes. Insoluble material was removed by centrifugation at 15,000 × g for 15 min at 4°C. Samples were then equalized using the Bradford protein assay (Biorad), and incubated with the indicated antibody. Immunoprecipitations were performed by gentle end over end rotation for 3 hours, followed by incubation with protein A/G-sepharose beads for 45 minutes, and then immune complexes were washed four times in cold RIPA buffer and resuspended in Laemmli loading buffer.

### Protein half-life determination

Cells were incubated with 200 μg/mL of cyclohexamide (Sigma-Aldrich) for the indicated times and lysed in Laemmli buffer loading buffer. Protein levels were detected by immunoblotting.

### Cell Proliferation and Clonogenic Assays

Cell numbers were assessed indirectly by using the sulforhodamine B (SRB) assay as described in [84].

Clonogenic assays were performed by seeding 100-1,000 cells/well in a 6 well plate and treating as indicated. Cells were allowed to recover and grow for 14-16 days. Cells were then washed twice with PBS and fixed and dyed with a solution containing: 0.5% crystal violet, 0.5% formal saline, 50% ethanol, and 145 mM NaCl for 20 minutes at room temperature. Cells were then washed three times in ddH2O and air dried overnight. Plates were scanned and colonies were manually counted in ImageJ with the CellCounter plugin.

### RNA isolation, reverse transcription and quantitative real time PCR (RT-qPCR) analysis

Total RNA was isolated from cells after the indicated treatments using the EZ-10 DNAaway RNA Miniprep kit (Bio Basic). Reverse transcription to generate cDNAs was performed using the qScript cDNA SuperMix (Quantabio). Quantitative real-time PCR analysis was performed using PerfeCTa SYBR Supermix with 20 ng of cDNA per reaction and 200 nM of the following specific primers:

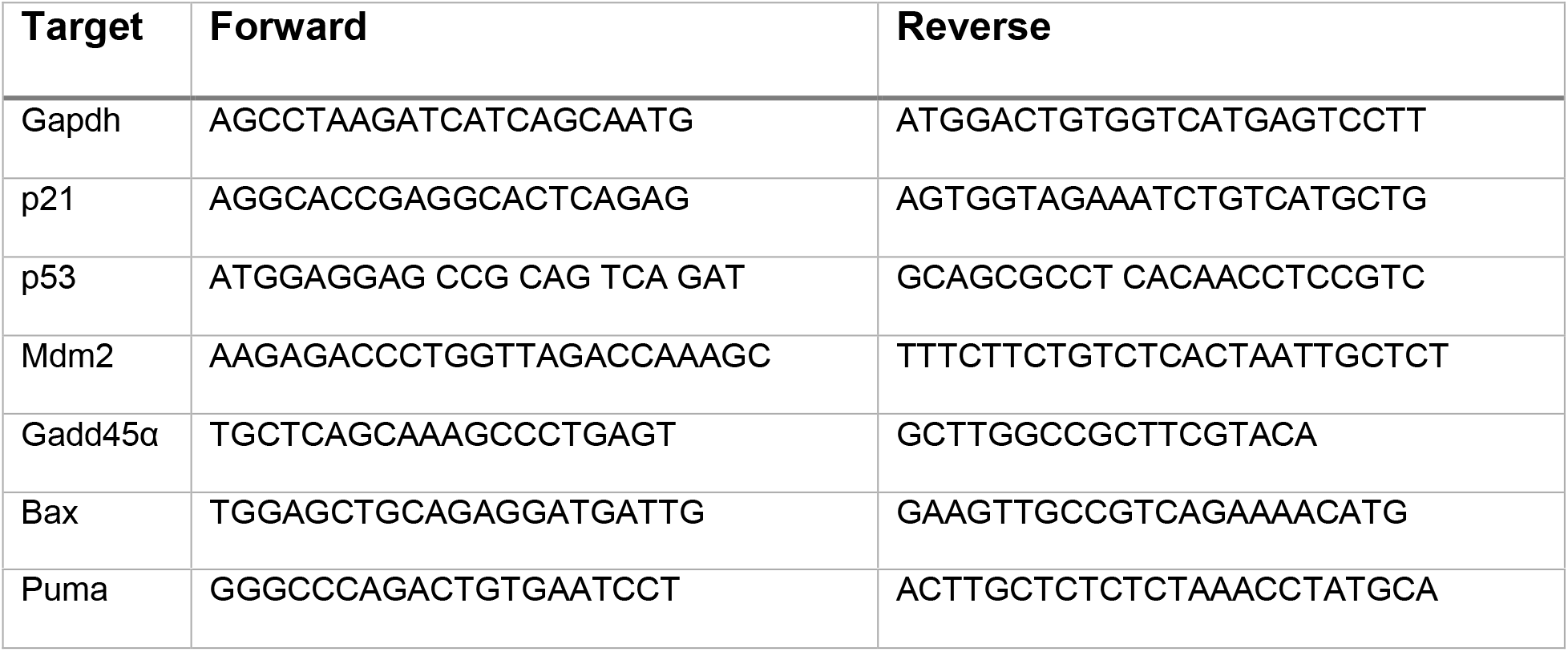

Quantitative real-time PCR was performed on the Roche LightCycler 480, and relative mRNA expression was calculated using the 2 ^−ΔΔCt^ method using GAPDH as a reference gene [85].

### Cell cycle analysis by flow cytometry

For both methods of cell cycle analysis, cells were seeded into 6 cm plates, treated as indicated, trypsinized, and fixed with ice cold 70% ethanol (−20°C) with gentle vortexing. Cells were then allowed to fix on ice for one hour.

For total DNA content analysis for cell cycle distribution, samples were washed twice with PBS and stained with 50 μg/mL propidium iodide and 20 μg/mL RNAse A in PBS + 0.1% Triton X-100. Stained cells were analysed on a FACSCanto flow cytometer (BD Biosciences) and CellQuest software (BD Immunocytometry Systems). 10^4^ events were analyzed for each sample on FlowJo.

For the quantification of mitotic cells, indirect immunofluorescence was used on ethanol fixed cells using an anti-phospho–H3 (Ser10) antibody (Millipore). Briefly, cells were washed twice with PBS, and permeabilized with 0.25% Triton X-100 and incubated on ice for 15 minutes. Cells were washed twice with PBS + 1% BSA and incubated with 0.25 μg anti-phospho–H3 (Ser10) antibody with 1% BSA for 1 hour at room temperature. Cells were then incubated in goat anti-mouse IgG conjugated to Alexa488 (1:300 dilution) with 1% BSA at room temperature for 30 minutes under light protection. Cells were then counterstained with a DNA content stain consisting of 50 μg/mL propidium iodide and 20 μg/mL RNAse A in PBS. Stained cells were analysed on a FACSCanto flow cytometer (BD Biosciences) and CellQuest software (BD Immunocytometry Systems). Twenty thousand events were analyzed for each sample on FlowJo.

### Annexin V/Propidium Iodide assay for apoptosis

For Annexin V/PI assays, cells were evaluated for apoptosis by flow cytometry according to the manufacturer’s protocol (BD Pharmingen, San Diego, CA, USA). Briefly, cells were trypsinized, washed with PBS and 2×10^5^ cells were stained with 5 μL of Annexin V-FITC and 10 μL of 50 ng/mL of propidium iodide in 200 μL binding buffer (10 mM HEPES, pH 7.4, 140 mM NaCl, 2.5 mM CaCl2) for 15 minutes at room temperature in the dark. The apoptotic cells were determined using a FACSCanto flow cytometer (BD Biosciences) and CellQuest software (BD Immunocytometry Systems). Ten thousand events were analyzed for each sample on FlowJo.

### Bacterial protein purification

GST-p53 and GST-p53 Y327F were produced in *BL21 E.coli* cells. Single colonies of transformed bacteria were inoculated as starter cultures and grown overnight at 37°C with gentle agitation. Starter cultures were diluted into large scale production cultures to an OD_600_ of 0.05-0.1. Production cultures were allowed to grow until an OD_600_ of 0.5 had been reached, subsequent to which cultures were incubated with 0.5 mM of IPTG for 4 hours. Cells were pelleted at 6000xg for 15 min, followed by snap freezing in liquid nitrogen. Cells were then thawed in a 37°C water bath and lysed with PBS +1% Triton X-100 + lysozyme. Cells were sonicated in 15 second pulses, followed by 15 seconds on ice three times, and then left on ice for 30 minutes. Insoluble material was removed via centrifugation at 15,000xg for 15 minutes. The supernatant was then incubated with 20 μL/mL of glutathione sepharose 4B resin for 2 hours. Beads were then washed three times with lysis buffer, and incubated with elution buffer (10 mM reduced glutathione, 50 mM Tris-HCl pH 7.6, 150 mM NaCl) three times for 10 mines. Eluted fractions were pooled and protein amount and purity was determined via SDS-PAGE Coomassie staining.

### *In Vitro* Kinase Assays

Kinase assays were performed by the incubation of 100-200 ng of purified NEK10 at 30°C for 30 minutes in kinase assay buffer supplemented with 5 μCi [γ-^32^P] ATP, 20 μM ATP and 1 μg of GST-p53 or GST-p53 Y327F. Reactions were terminated using electrophoresis buffer followed by boiling for 5 minutes. Samples were resolved with SDS-PAGE, and imaged on a Typhoon Imager (GE Healthcare Lifesciences). Relative activity was determined using densitometry on ImageJ.

### EdU Pulse Labelling for DNA synthesis determination

To determine the proportion of cells with active DNA synthesis, we used a modified EdU pulse labeling protocol that utilizes Click chemistry. Briefly, cells were grown and pulse labeled with 10 μM EdU (Setareh Biotech) for 2 hours at 37°C with 5% CO2. Cells were then trypsinized and fixed with 4% paraformaldehyde, permeabilized with 0.1% Triton X-100, washed with PBS +1% BSA. Permeabilized cells were dyed with a Click labelling dye (8 μM FAM-Azide 488 (Lumiprobe), 2 mM CuSO4, 20 mg/mL ascorbic acid) for 30 minutes in the dark at room temperature. 10,000 cells were counted and analysed using a FACSCanto flow cytometer (BD Biosciences) and CellQuest software (BD Immunocytometry Systems).

### Oligomerization Assay

Cells were treated as indicated and lysed with a buffer containing: 25 mM Hepes pH 7.5, 150 mM NaCl, and 1% NP-40, supplemented with fresh 1 mM dithiothreitol (DTT), 0.1 mM sodium orthovanadate and a protease inhibitor cocktail (Sigma). Lysates were incubated on ice for 20 min, and insoluble material was removed by centrifugation at 15,000 × g for 15 min at 4°C. Samples were then equalized using the Bradford protein assay, and treated with 0.025% Glutaraldehyde for 30 minutes on ice. The glutaraldehyde was quenched with the addition of 100 mM Glycine pH 2.5.

### Bioinformatics Analysis

RNASeq V2 and clinical data from The Cancer Genome Atlas (TCGA) Breast datasets downloaded from the UCSC XENA platform were used to generate Kaplan-Meier survival curves (xena.ucsc.edu) [86]. Median expression of *NEK10* was used to divide the “*NEK10* High” and “*NEK10* Low” groups.

## SUPPLEMENTAL FIGURES

**Figure S1.**
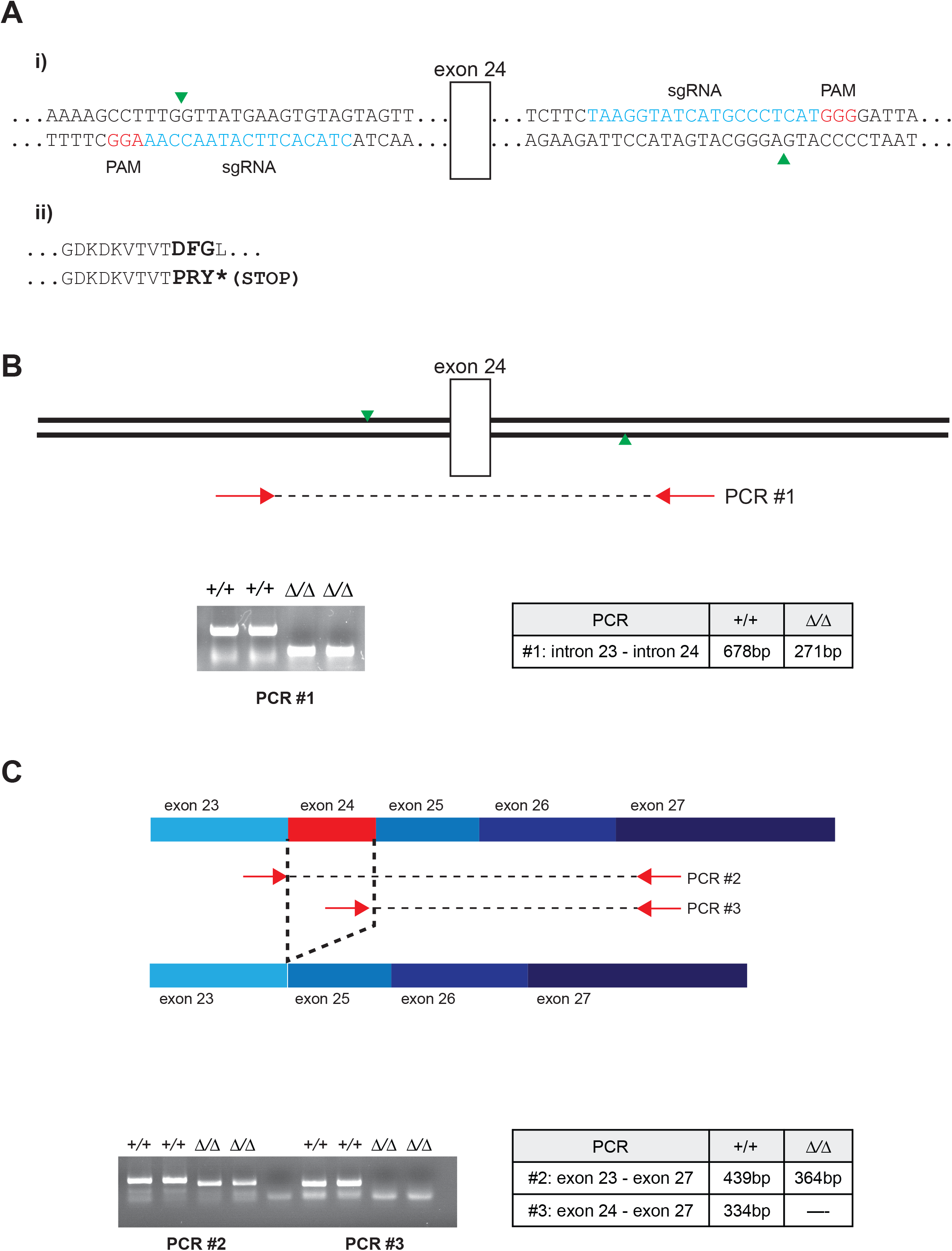
CRISPR-Cas9-mediated *NEK10* deletion strategy. **A) i**. Deletion of *NEK10* exon 24, encoding the critical DFG kinase domain motif, was engineered using paired sgRNA-guided Cas9 nickases to introduce cuts in intronic sequences upstream and downstream of the targeted exon. The specific sequences targeted by the sgRNAs are represented in blue and the predicted cut sites by green arrowheads. **ii**. The loss of exon 24 generates a frameshift mutation that introduces a stop codon 3aa downstream of exon 23-encoded sequence. **B**) Exon 24 deletion was initially established by means of genomic PCR using primers (red arrows) external to the predicted cut sites (green arrowheads). The predicted PCR product sizes and a representative gel are shown. **C**) Two RT-PCR reactions were utilized as additional confirmation of exon 24 deletion. The first reaction set (PCR#2) employed primers (red arrows) upstream and downstream of the targeted exon such that the loss of exon 24 could be detected by a reduction in the size of the PCR product. The second reaction set (PCR#3) employed an upstream primer within the deleted exon enabling the loss of exon 24 to be established by the absence of PCR product. The predicted PCR product sizes and a representative gel are shown.

**Figure S2.**
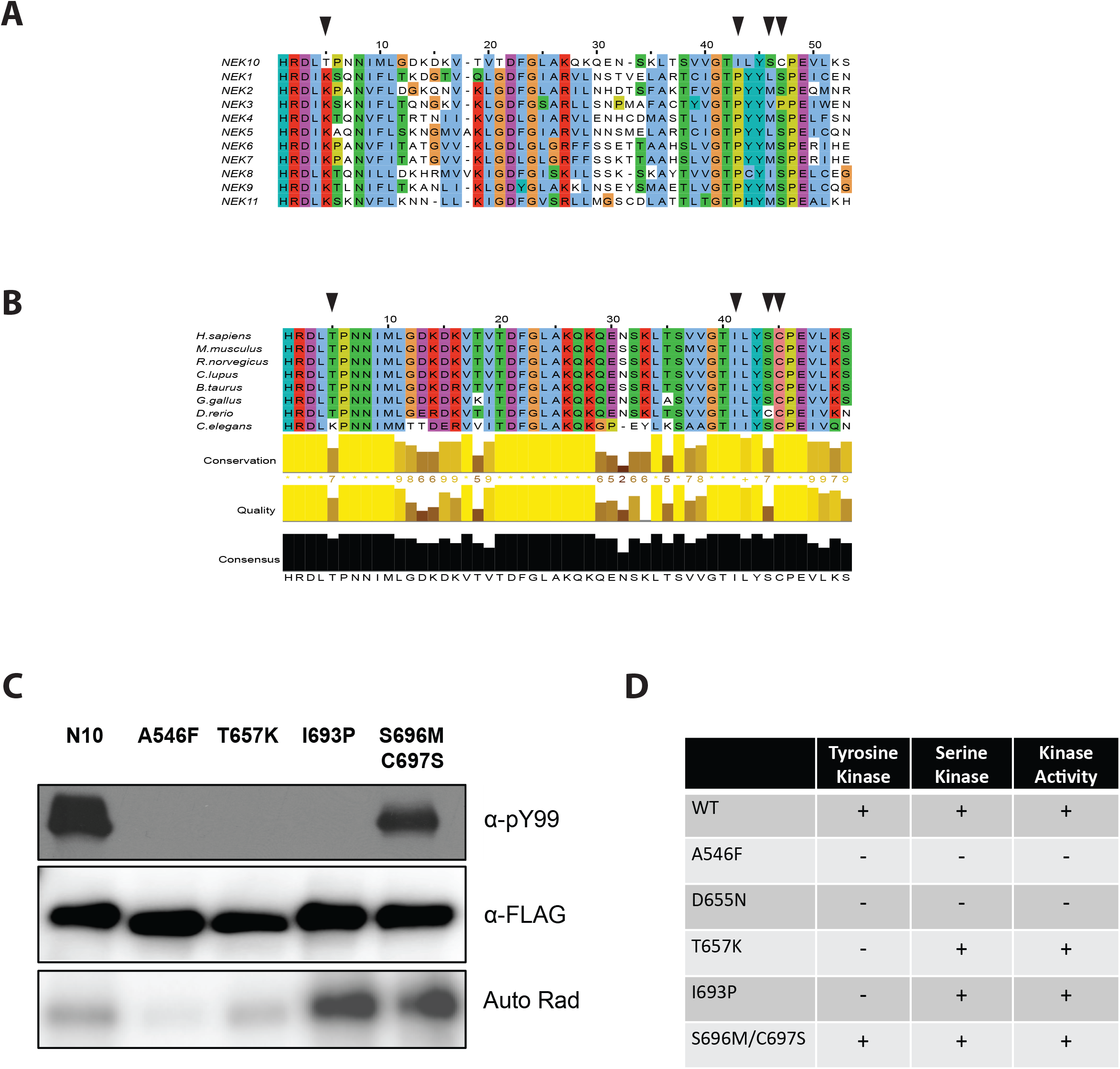
I693 in the activation loop is required for tyrosine kinase activity. **A**) The catalytic and activation loops of NEK kinase family members were aligned with CLUSTAL Omega, residues unique to NEK10 are indicated with arrows. **B**) The catalytic and activation loops of NEK10 homologues from the indicated species were aligned using CLUSTAL Omega to determine the degree of conservation of unique amino acids. **C**) Purified NEK10 and activation loop mutants were analysed for their ability to autophosphorylate on tyrosines via radioactive kinase assay for the total kinase activity and via western blot for tyrosine activity. **D**) Summary table indicating substrate specificity for NEK10 mutants.

**Figure S3.**
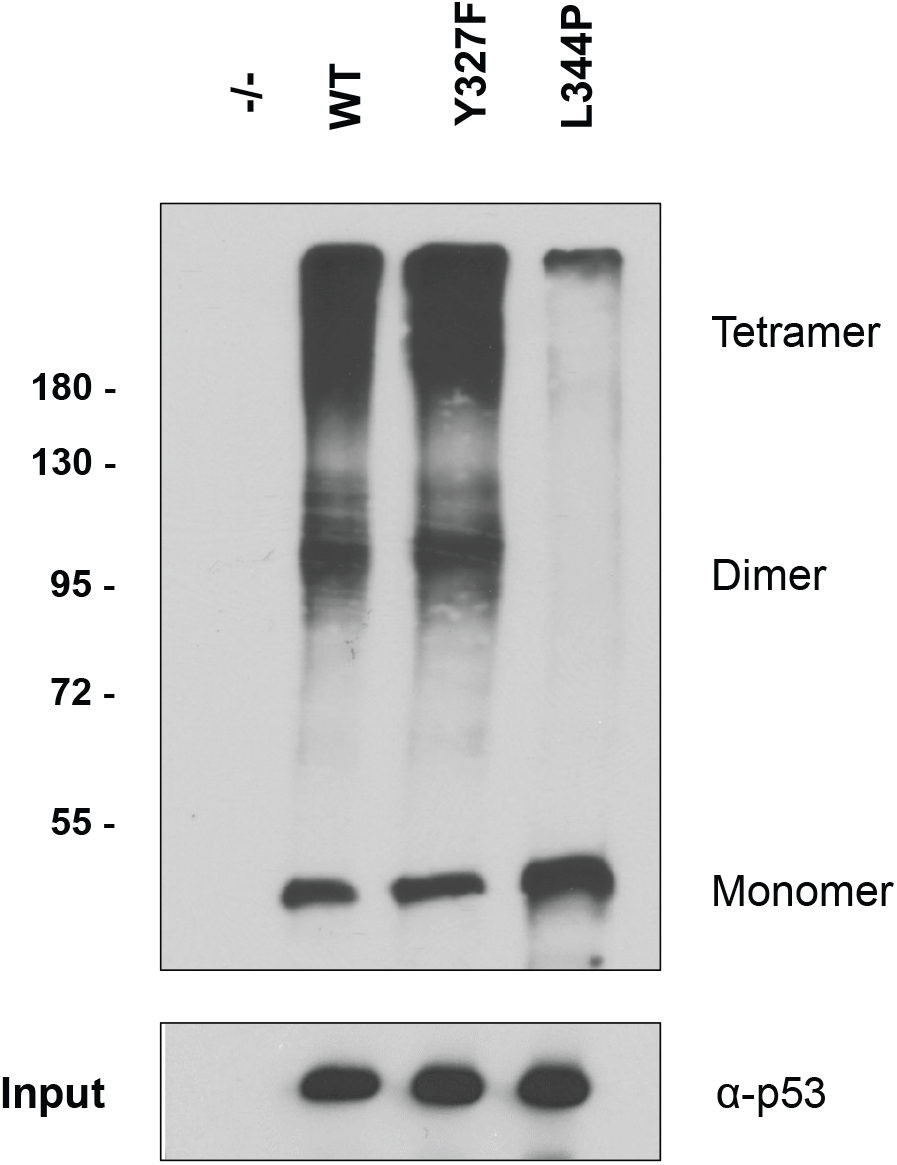
Y327F mutation does not affect p53 oligomerization. HCT116 *p53^−/−^* cells expressing the indicated constructs were lysed and lysates were treated with 0.025% glutaraldehyde to cross link proteins and immunoblotted to assess the degree of p53 oligomerization.

## References

1. Menendez, D., A. Inga, and M.A. Resnick, The expanding universe of p53 targets. Nat Rev Cancer, 2009. 9(10): p. 724–37.

2. Kastenhuber, E.R. and S.W. Lowe, Putting p53 in Context. Cell, 2017. 170(6): p. 1062–1078.

3. Horn, H.F. and K.H. Vousden, Coping with stress: multiple ways to activate p53. Oncogene, 2007. 26(9): p. 1306–16.

4. Pappas, K., et al., p53 Maintains Baseline Expression of Multiple Tumor Suppressor Genes. Mol Cancer Res, 2017. 15(8): p. 1051–1062.

5. Allen, M.A., et al., Global analysis of p53-regulated transcription identifies its direct targets and unexpected regulatory mechanisms. Elife, 2014. 3: p. e02200.

6. Fischer, M., Census and evaluation of p53 target genes. Oncogene, 2017. 36(28): p. 3943–3956.

7. McGranahan, N. and C. Swanton, Clonal Heterogeneity and Tumor Evolution: Past, Present, and the Future. Cell, 2017. 168(4): p. 613–628.

8. Malkin, D., Li-fraumeni syndrome. Genes Cancer, 2011. 2(4): p. 475–84.

9. Riley, T., et al., Transcriptional control of human p53-regulated genes. Nat Rev Mol Cell Biol, 2008. 9(5): p. 402–12.

10. Huang, L.C., K.C. Clarkin, and G.M. Wahl, Sensitivity and selectivity of the DNA damage sensor responsible for activating p53-dependent G1 arrest. Proc Natl Acad Sci U S A, 1996. 93(10): p. 4827–32.

11. Kumari, R., S. Kohli, and S. Das, p53 regulation upon genotoxic stress: intricacies and complexities. Mol Cell Oncol, 2014. 1(3): p. e969653.

12. Haupt, Y., et al., Mdm2 promotes the rapid degradation of p53. Nature, 1997. 387(6630): p. 296–9.

13. Brooks, C.L. and W. Gu, p53 ubiquitination: Mdm2 and beyond. Mol Cell, 2006. 21(3): p. 307–15.

14. Canman, C.E., et al., Activation of the ATM kinase by ionizing radiation and phosphorylation of p53. Science, 1998. 281(5383): p. 1677–9.

15. Stommel, J.M. and G.M. Wahl, Accelerated MDM2 auto-degradation induced by DNA-damage kinases is required for p53 activation. EMBO J, 2004. 23(7): p. 1547–56.

16. Tibbetts, R.S., et al., A role for ATR in the DNA damage-induced phosphorylation of p53. Genes Dev, 1999. 13(2): p. 152–7.

17. Schon, O., et al., Molecular mechanism of the interaction between MDM2 and p53. J Mol Biol, 2002. 323(3): p. 491–501.

18. Banin, S., et al., Enhanced phosphorylation of p53 by ATM in response to DNA damage. Science, 1998. 281(5383): p. 1674–7.

19. Tang, Y., et al., Acetylation is indispensable for p53 activation. Cell, 2008. 133(4): p. 612–26.

20. Sykes, S.M., et al., Acetylation of the p53 DNA-binding domain regulates apoptosis induction. Mol Cell, 2006. 24(6): p. 841–51.

21. Tang, Y., et al., Tip60-dependent acetylation of p53 modulates the decision between cell-cycle arrest and apoptosis. Mol Cell, 2006. 24(6): p. 827–39.

22. Sykes, S.M., et al., Acetylation of the DNA binding domain regulates transcription-independent apoptosis by p53. J Biol Chem, 2009. 284(30): p. 20197–205.

23. Chao, C., et al., Acetylation of mouse p53 at lysine 317 negatively regulates p53 apoptotic activities after DNA damage. Mol Cell Biol, 2006. 26(18): p. 6859–69.

24. Liu, L., et al., p53 sites acetylated in vitro by PCAF and p300 are acetylated in vivo in response to DNA damage. Mol Cell Biol, 1999. 19(2): p. 1202–9.

25. Knights, C.D., et al., Distinct p53 acetylation cassettes differentially influence gene-expression patterns and cell fate. J Cell Biol, 2006. 173(4): p. 533–44.

26. Jansson, M., et al., Arginine methylation regulates the p53 response. Nat Cell Biol, 2008. 10(12): p. 1431–9.

27. Chen, Y., et al., Never-in-mitosis related kinase 1 functions in DNA damage response and checkpoint control. Cell Cycle, 2008. 7(20): p. 3194–201.

28. Chen, Y., et al., Nek1 kinase functions in DNA damage response and checkpoint control through a pathway independent of ATM and ATR. Cell Cycle, 2011. 10(4): p. 655–63.

29. Sabir, S.R., et al., Loss of Nek11 Prevents G2/M Arrest and Promotes Cell Death in HCT116 Colorectal Cancer Cells Exposed to Therapeutic DNA Damaging Agents. PLoS One, 2015. 10(10): p. e0140975.

30. Melixetian, M., et al., NEK11 regulates CDC25A degradation and the IR-induced G2/M checkpoint. Nat Cell Biol, 2009. 11(10): p. 1247–53.

31. Noguchi, K., et al., Nek11, a new member of the NIMA family of kinases, involved in DNA replication and genotoxic stress responses. J Biol Chem, 2002. 277(42): p. 39655–65.

32. Fletcher, L., et al., Inhibition of centrosome separation after DNA damage: a role for Nek2. Radiat Res, 2004. 162(2): p. 128–35.

33. Mi, J., et al., Protein phosphatase-1alpha regulates centrosome splitting through Nek2. Cancer Res, 2007. 67(3): p. 1082–9.

34. Lee, M.Y., et al., Nek6 is involved in G2/M phase cell cycle arrest through DNA damage-induced phosphorylation. Cell Cycle, 2008. 7(17): p. 2705–9.

35. Choi, H.J., et al., NEK8 links the ATR-regulated replication stress response and S phase CDK activity to renal ciliopathies. Mol Cell, 2013. 51(4): p. 423–39.

36. Abeyta, A., et al., NEK8 regulates DNA damage-induced RAD51 foci formation and replication fork protection. Cell Cycle, 2017. 16(4): p. 335–347.

37. Smith, S.C., et al., A gemcitabine sensitivity screen identifies a role for NEK9 in the replication stress response. Nucleic Acids Res, 2014. 42(18): p. 11517–27.

38. Moniz, L.S. and V. Stambolic, Nek10 mediates G2/M cell cycle arrest and MEK autoactivation in response to UV irradiation. Mol Cell Biol, 2011. 31(1): p. 30–42.

39. Forbes, S.A., et al., COSMIC: somatic cancer genetics at high-resolution. Nucleic Acids Res, 2017. 45(D1): p. D777–D783.

40. Gao, J., et al., Integrative analysis of complex cancer genomics and clinical profiles using the cBioPortal. Sci Signal, 2013. 6(269): p. pl1.

41. Cerami, E., et al., The cBio cancer genomics portal: an open platform for exploring multidimensional cancer genomics data. Cancer Discov, 2012. 2(5): p. 401–4.

42. Ahmed, S., et al., Newly discovered breast cancer susceptibility loci on 3p24 and 17q23.2. Nat Genet, 2009. 41(5): p. 585–90.

43. Mendoza, M.C., E.E. Er, and J. Blenis, The Ras-ERK and PI3K-mTOR pathways: cross-talk and compensation. Trends Biochem Sci, 2011. 36(6): p. 320–8.

44. Cazzalini, O., et al., Multiple roles of the cell cycle inhibitor p21(CDKN1A) in the DNA damage response. Mutat Res, 2010. 704(1-3): p. 12–20.

45. Besson, A., S.F. Dowdy, and J.M. Roberts, CDK inhibitors: cell cycle regulators and beyond. Dev Cell, 2008. 14(2): p. 159–69.

46. Giono, L.E. and J.J. Manfredi, The p53 tumor suppressor participates in multiple cell cycle checkpoints. J Cell Physiol, 2006. 209(1): p. 13–20.

47. Junttila, M.R. and G.I. Evan, p53--a Jack of all trades but master of none. Nat Rev Cancer, 2009. 9(11): p. 821–9.

48. Moniz, L.S., Characterization of NimA-related kinase 10 (NEK10): A role in checkpoint control. 2010, University of Toronto, 2010. p. 178 leaves.

49. Huang, Y.F. and D.V. Bulavin, Oncogene-mediated regulation of p53 ISGylation and functions. Oncotarget, 2014. 5(14): p. 5808–18.

50. Tsai, C.F., et al., Large-scale determination of absolute phosphorylation stoichiometries in human cells by motif-targeting quantitative proteomics. Nat Commun, 2015. 6: p. 6622.

51. Bai, Y., et al., Phosphoproteomics identifies driver tyrosine kinases in sarcoma cell lines and tumors. Cancer Res, 2012. 72(10): p. 2501–11.

52. Kawaguchi, T., et al., The relationship among p53 oligomer formation, structure and transcriptional activity using a comprehensive missense mutation library. Oncogene, 2005. 24(46): p. 6976–81.

53. Yakovlev, V.A., et al., Nitration of the tumor suppressor protein p53 at tyrosine 327 promotes p53 oligomerization and activation. Biochemistry, 2010. 49(25): p. 5331–9.

54. Chong, L.T., et al., Kinetic computational alanine scanning: application to p53 oligomerization. J Mol Biol, 2006. 357(3): p. 1039–49.

55. Davies, H., et al., Somatic mutations of the protein kinase gene family in human lung cancer. Cancer Res, 2005. 65(17): p. 7591–5.

56. Greenman, C., et al., Patterns of somatic mutation in human cancer genomes. Nature, 2007. 446(7132): p. 153–8.

57. Ebctcg, et al., Effect of radiotherapy after mastectomy and axillary surgery on 10-year recurrence and 20-year breast cancer mortality: meta-analysis of individual patient data for 8135 women in 22 randomised trials. Lancet, 2014. 383(9935): p. 2127–35.

58. Early Breast Cancer Trialists’ Collaborative, G., et al., Effect of radiotherapy after breast-conserving surgery on 10-year recurrence and 15-year breast cancer death: meta-analysis of individual patient data for 10,801 women in 17 randomised trials. Lancet, 2011. 378(9804): p. 1707–16.

59. Lakin, N.D. and S.P. Jackson, Regulation of p53 in response to DNA damage. Oncogene, 1999. 18(53): p. 7644–55.

60. Meek, D.W., The p53 response to DNA damage. DNA Repair (Amst), 2004. 3(8-9): p. 1049–56.

61. Jee, H.J., et al., Nek6 overexpression antagonizes p53-induced senescence in human cancer cells. Cell Cycle, 2010. 9(23): p. 4703–10.

62. Salem, H., et al., Nek7 kinase targeting leads to early mortality, cytokinesis disturbance and polyploidy. Oncogene, 2010. 29(28): p. 4046–57.

63. Moniz, L., et al., Nek family of kinases in cell cycle, checkpoint control and cancer. Cell Div, 2011. 6: p. 18.

64. Wang, S. and W.S. El-Deiry, p73 or p53 directly regulates human p53 transcription to maintain cell cycle checkpoints. Cancer Res, 2006. 66(14): p. 6982–9.

65. Hudson, J.M., R. Frade, and M. Bar-Eli, Wild-type p53 regulates its own transcription in a cell-type specific manner. DNA Cell Biol, 1995. 14(9): p. 759–66.

66. Deffie, A., et al., The tumor suppressor p53 regulates its own transcription. Mol Cell Biol, 1993. 13(6): p. 3415–23.

67. Purvis, J.E., et al., p53 dynamics control cell fate. Science, 2012. 336(6087): p. 1440–4.

68. Chen, X., et al., DNA damage strength modulates a bimodal switch of p53 dynamics for cell-fate control. BMC Biol, 2013. 11: p. 73.

69. Stewart-Ornstein, J. and G. Lahav, p53 dynamics in response to DNA damage vary across cell lines and are shaped by efficiency of DNA repair and activity of the kinase ATM. Sci Signal, 2017. 10(476).

70. Espinosa, J.M., R.E. Verdun, and B.M. Emerson, p53 functions through stress-and promoter-specific recruitment of transcription initiation components before and after DNA damage. Mol Cell, 2003. 12(4): p. 1015–27.

71. Morachis, J.M., C.M. Murawsky, and B.M. Emerson, Regulation of the p53 transcriptional response by structurally diverse core promoters. Genes Dev, 2010. 24(2): p. 135–47.

72. Batchelor, E., et al., Stimulus-dependent dynamics of p53 in single cells. Mol Syst Biol, 2011. 7: p. 488.

73. Brady, C.A., et al., Distinct p53 transcriptional programs dictate acute DNA-damage responses and tumor suppression. Cell, 2011. 145(4): p. 571–83.

74. Jin, S., et al., GADD45-induced cell cycle G2-M arrest associates with altered subcellular distribution of cyclin B1 and is independent of p38 kinase activity. Oncogene, 2002. 21(57): p. 8696–704.

75. Wang, X.W., et al., GADD45 induction of a G2/M cell cycle checkpoint. Proc Natl Acad Sci U S A, 1999. 96(7): p. 3706–11.

76. Taylor, W.R. and G.R. Stark, Regulation of the G2/M transition by p53. Oncogene, 2001. 20(15): p. 1803–15.

77. Lord, C.J. and A. Ashworth, BRCAness revisited. Nat Rev Cancer, 2016. 16(2): p. 110–20.

78. Tutt, A., et al., Carboplatin in BRCA 1/2-mutated and triple-negative breast cancer BRCAness subgroups: the TNT Trial. Nat Med, 2018. 24(5): p. 628–637.

79. De Summa, S., et al., BRCAness: a deeper insight into basal-like breast tumors. Ann Oncol, 2013. 24 Suppl 8: p. viii13–viii21.

80. Shah, S.P., et al., The clonal and mutational evolution spectrum of primary triple-negative breast cancers. Nature, 2012. 486(7403): p. 395–9.

81. Curtis, C., et al., The genomic and transcriptomic architecture of 2,000 breast tumours reveals novel subgroups. Nature, 2012. 486(7403): p. 346–52.

82. Burstein, M.D., et al., Comprehensive genomic analysis identifies novel subtypes and targets of triple-negative breast cancer. Clin Cancer Res, 2015. 21(7): p. 1688–98.

83. Lehmann, B.D., et al., Refinement of Triple-Negative Breast Cancer Molecular Subtypes: Implications for Neoadjuvant Chemotherapy Selection. PLoS One, 2016. 11(6): p. e0157368.

84. Vichai, V. and K. Kirtikara, Sulforhodamine B colorimetric assay for cytotoxicity screening. Nat Protoc, 2006. 1(3): p. 1112–6.

85. Schmittgen, T.D. and K.J. Livak, Analyzing real-time PCR data by the comparative C(T) method. Nat Protoc, 2008. 3(6): p. 1101–8.

86. Goldman, M., et al., The UCSC Xena Platform for cancer genomics data visualization and interpretation. bioRxiv, 2018.

